# HiFi chromosome-scale diploid assemblies of the grape rootstocks 110R, Kober 5BB, and 101-14 Mgt

**DOI:** 10.1101/2022.07.29.502067

**Authors:** Andrea Minio, Noé Cochetel, Mélanie Massonnet, Rosa Figueroa-Balderas, Dario Cantu

## Abstract

Cultivated grapevines are commonly grafted on closely related species to cope with specific biotic and abiotic stress conditions. The three North American *Vitis* species *V. riparia*, *V. rupestris*, and *V. berlandieri*, are the main species used for breeding grape rootstocks. Here, we report the diploid chromosome-scale assembly of three widely used rootstocks derived from these species: Richter 110 (110R), Kober 5BB, and 101-14 Millardet et de Grasset (Mgt). Draft genomes of the three hybrids were assembled using PacBio HiFi sequences at an average coverage of 53.1 X-fold. Using the tool suite HaploSync, we reconstructed the two sets of nineteen chromosome-scale pseudomolecules for each genome with an average haploid genome size of 494.5 Mbp. Residual haplotype switches were resolved using shared-haplotype information. These three reference genomes represent a valuable resource for studying the genetic basis of grape adaption to biotic and abiotic stresses, and designing trait-associated markers for rootstock breeding programs.

## Background & Summary

Cultivated grapevines (*Vitis vinifera* ssp. *vinifera*) are usually grafted onto rootstocks derived from North American *Vitis* species (Figure 1A). This practice was established during the 19th century in response to the near devastation of European vineyards by the grape root aphid phylloxera (*Daktulosphaira vitifoliae* Fitch)^1^. Grape phylloxera was introduced into Europe in the 1850s through the movement of plant material from North America^2^. Most North American *Vitis* species are resistant to phylloxera, likely as a result of co-evolution with the insect in their native environment. *Vitis riparia* and *V. rupestris* were the first wild grape species used as rootstock because they root easily from hardwood cuttings and have good grafting compatibility with the berry-producing scions^3^. However, these two species were not suitable for calcareous soils, which are common in Europe. *Vitis berlandieri*, another North American grape species, was then found to be resistant to phylloxera and lime-tolerant, although it poorly roots from dormant cuttings^4^. To introduce the lime-tolerance of *V. berlandieri* and improve its rootability, new rootstocks were bred crossing *V. berlandieri* with either *V. riparia* or *V. rupestris*. Today, commercialized rootstocks are mainly hybrids of these three grape species^5^. Among these, Richter 110 (110R; *V. berlandieri* x *V. rupestris*), Kober 5BB (*V. berlandieri* x *V. riparia*), and 101-14 Millardet et de Grasset (Mgt; *V. riparia* x *V. rupestris*) are the most commonly used worldwide. In addition to their resistance to phylloxera, grape rootstocks are choosen based on tolerance to biotic (e.g. nematodes) and abiotic stresses (e.g. drought), preference of soil physicochemical properties, and the vigor level they confer to the scion^6^. For instance, 101-14 Mgt generally triggers the precocity of the vegetative growth despite a moderate vigor, whereas 110R and Kober 5BB confer high vigor and delay plant maturity^7^. 110R is known for its drought tolerance and excess soil moisture has negative impacts on its development^6^. In contrast, 101-14 Mgt and Kober 5BB are not considered drought-tolerant and grow well in moist soils^6^. The three rootstocks also have different levels of tolerance to nematodes depending on the nematode species^6, 8^.

**Figure 1:**
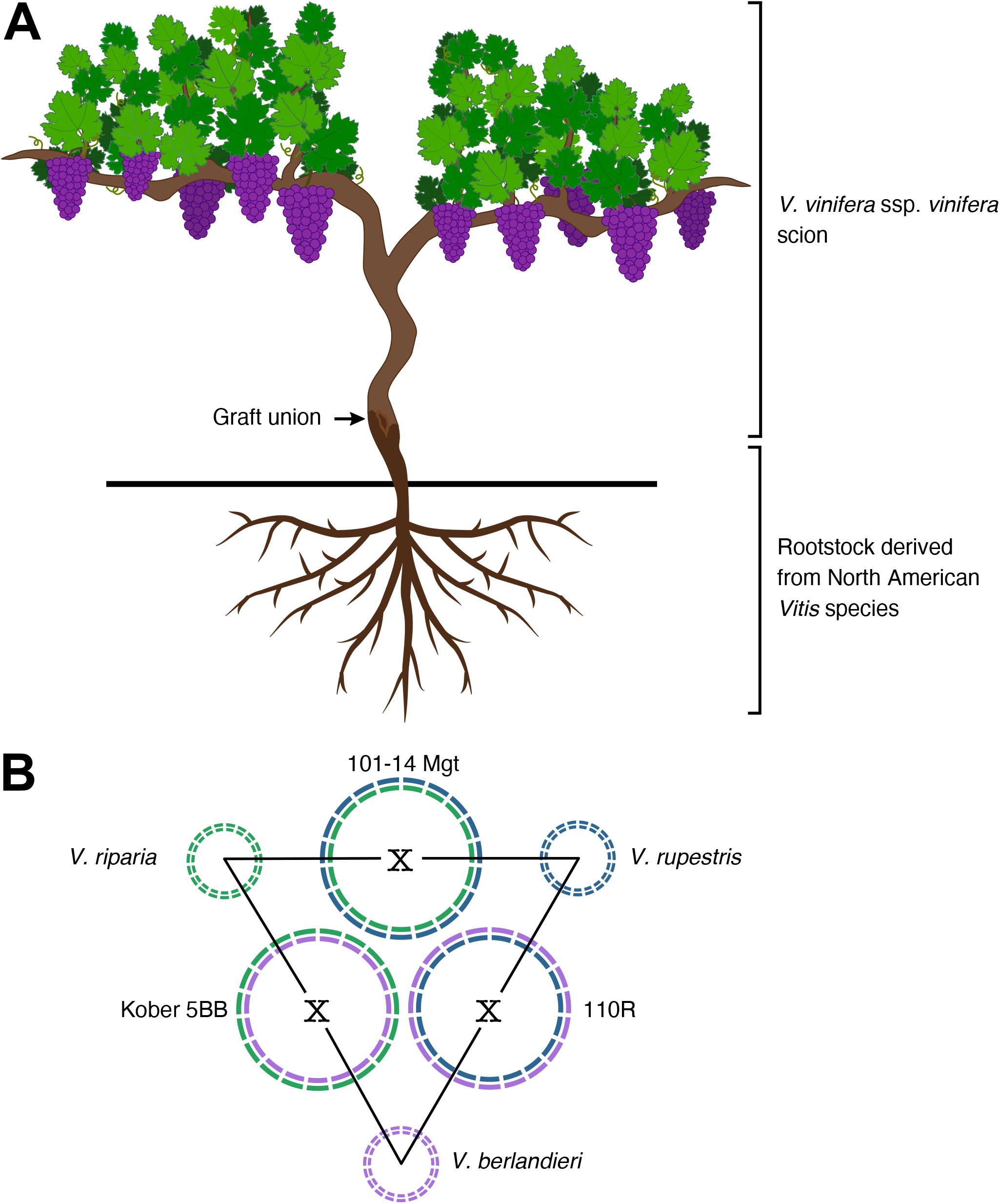
Description of the three grape rootstocks 101-14 Mgt, 110R, and Kober 5BB. A) Wine grapevine scion (*Vitis vinifera* spp. *vinifera*) grafted onto a rootstock from another *Vitis* species. B) Schematic representation of haplotype composition of 101-14 Mgt, 110R, and Kober 5BB. Each pair of rootstocks shares a set of chromosomes from the same parental *Vitis* species. Shared haplotypes are represented with the same color.

In addition to their commercial importance, rootstocks are valuable to study the genetic bases of grape adaptation to biotic and abiotic stresses^9^. However, to date only two genomes of *V. riparia* have been published^10, 11^ and no genome reference is available for any of the commonly used rootstocks. This article describes the chromosome-scale diploid genome assembly of 110R, Kober 5BB, and 101-14 Mgt. Genomes were sequenced using highly accurate long-read sequencing (HiFi, Pacific Biosciences) and assembled with Hifiasm^12^. Each diploid draft genome was then scaffolded into two sets of pseudomolecules using the tool suite HaploSync^13^, and haplotypes were assigned to each *Vitis* parent based on sequence similarity between the haplotypes derived from the same species. These genomes represent an important resource for investigating the genetic basis of resistance to environmental factors and designing markers to accelerate rootstock breeding programs.

## Methods

### Library preparation and sequencing

Young leaves (1-2 cm-wide) were collected from 110R (FPS 01), Kober 5BB (FPS 06), and 101-14 Mgt (FPS 01) at Foundation Plant Services (University of California Davis, Davis, CA) and immediately frozen and ground to powder in liquid nitrogen. High molecular weight genomic DNA was extracted from 1g of ground leaf tissue as described in Chin et al. (2016)^14^, and 12 μg of high molecular weight gDNA was sheared to a size distribution between 15 and 20 kbp using the Megaruptor^®^ 2 (Diagenode, Denville, NJ, USA). For each accession, one HiFi sequencing library was prepared using the SMRTbell^™^ Express Template Prep Kit 2.0 followed by immediate treatment with the Enzyme Clean Up Kit (Pacific Biosciences, Menlo Park, CA, USA). Libraries were size-selected using a BluePippin (Sage Sciences, Beverly, MA, USA) and HiFi SMRTbell templates longer than 15 kbp were collected. Size-selected library fractions were cleaned using AMPure PB beads (Pacific Biosciences, Menlo Park, CA, USA). Concentration and final size distribution of the libraries were evaluated using a Qubit^™^ 1X dsDNA HS Assay Kit (Thermo Fisher, Waltham, MA, USA) and Femto Pulse System (Agilent, Santa Clara, CA, USA), respectively. HiFi libraries of 110R and Kober 5BB were sequenced using a PacBio Sequel II system (Pacific Biosciences, CA, USA) at the DNA Technology Core Facility, University of California, Davis (Davis, CA, USA). For 101-14 Mgt, sequencing was performed by Corteva Agriscience (Johnston, IA, USA) as an award from Pacific Biosciences to Dr. Noé Cochetel. An average of 26.5 ± 3.8 Gbp sequences were generated for each genome, corresponding to 53.1 ± 7.7 X-fold coverage of a 500 Mbp haploid genome (Table 1).

**Table 1:**
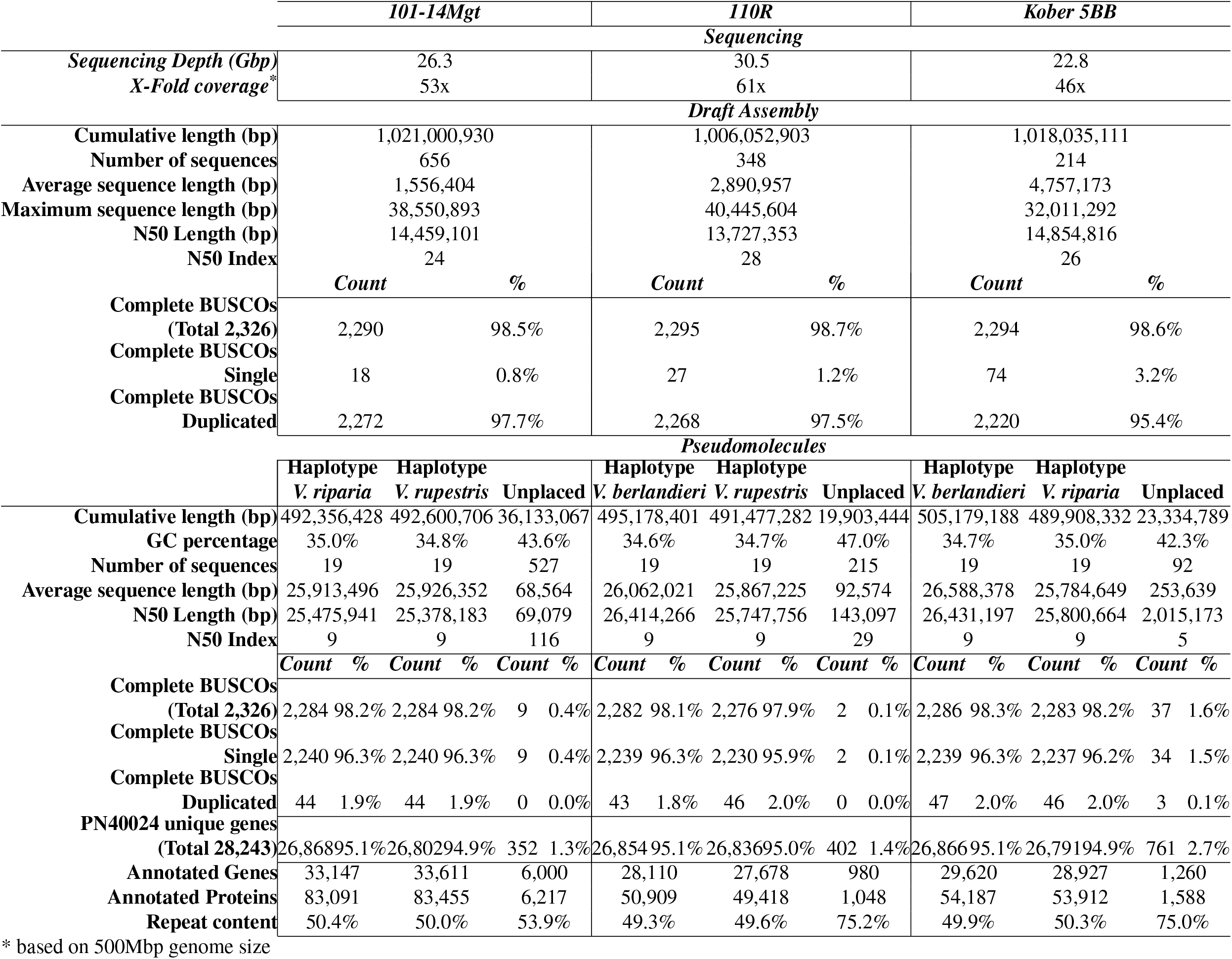
Genome assembly statistics of the three rootstocks. Summary statistics of the genome sequencing, draft genome assembly, chromosome-scale genome assembly, and gene annotation of 101-14 Mgt, 110R, and Kober 5BB rootstocks

Total RNA from *V. berlandieri* 9031, *V. rupestris* B38, and *V. riparia* 588271 leaves was isolated using a Cetyltrimethyl Ammonium Bromide (CTAB)-based extraction protocol as described in Blanco-Ulate et al. (2013)^15^. RNA purity was evaluated with a Nanodrop 2000 spectrophotometer (Thermo Scientific, Hanover Park, IL, USA), and RNA integrity by electrophoresis and an Agilent 2100 Bioanalyzer (Agilent Technologies, CA, USA). RNA quantity was assessed with a Qubit 2.0 Fluorometer and a broad range RNA kit (Life Technologies, Carlsbad, CA, USA). Total RNA (300 ng, RNA Integrity Number > 8.0) were used for library construction. Short-read cDNA libraries were prepared using the Illumina TruSeq RNA sample preparation kit v.2 (Illumina, CA, USA) following Illumina’s low-throughput protocol. Libraries were evaluated for quantity and quality with the High Sensitivity chip and an Agilent 2100 Bioanalyzer (Agilent Technologies, CA, USA). One library per species was sequenced using an Illumina HiSeq4000 sequencer with a 2×100bp protocol (DNA Technology Core Facility, University of California, Davis, CA, USA). Long-read cDNA SMRTbell libraries were prepared for *V. berlandieri* and *V. riparia*. First-strand synthesis and cDNA amplification were accomplished using the NEB Next Single Cell/Low Input cDNA Synthesis & Amplification Module (New England, Ipswich, MA, USA). The cDNAs were subsequently purified with ProNex magnetic beads (Promega, WI, USA) following the instructions in the Iso-Seq Express Template Preparation for Sequel and Sequel II Systems protocol (Pacific Biosciences, Menlo Park, CA, USA). ProNex magnetic beads (86 μL) were used to select amplified cDNA (≥ 2 kbp). At least 80 ng of the size-selected amplified cDNA were used to prepare the cDNA SMRTbell library. DNA damage repair and SMRTbell ligation was performed with SMRTbell Express Template Prep Kit 2.0 (Pacific Biosciences, Menlo Park, CA, USA) following the manufacturer’s protocol. One SMRT cell was sequenced for each species on the PacBio Sequel I platform (DNA Technology Core Facility, University of California, Davis, CA, USA).

### Genome assembly and pseudomolecule construction

HiFi reads were assembled using Hifiasm v.0.16.1-r374^12^. Multiple combinations of several assembly parameters were tested. A total of 1,939 assemblies were generated. The least fragmented assembly of each genotype was selected. The selected draft assemblies consisted of 406 ± 226 contigs with a N50 = 14.3 ± 0.6 Mbp (Table 1). Compared to other grape genomes previously generated with PacBio CLR technology, the PacBio HiFi reads greatly improves the contiguity of the draft assembly (PacBio CLR 1.2 ± 0.3 Mbp, Figure 2A). Gene space completeness was assessed using BUSCO V.5.1 with the Viridiplantae and Embryophyta ODB10 datasets^16^ and by mapping PN40024 (V1 annotation^17^) single-copy genes using GMAP v.2019-09-12 (alignments with at least 80% coverage and 80% identity were considered). For each rootstock, the draft genome assembly underwent quality control and scaffolding into a diploid set of chromosome-scale pseudomolecules using HaploSync^13^ and the *Vitis* consensus genetic map developed by Zou et al. (2020)^18^. One cycle of HaploFill was used for each genotype. The use of PacBio HiFi reads reduced significantly the fragmentation of the draft assembly compared to recently published grape genomes sequenced using PacBio CCS technology (Figure 2B)^13, 14, 19^. The lower fragmentation resulted in a 15 times smaller number of contigs necessary to scaffold a pseudomolecule (3.6 ± 2.0 HiFi contigs/pseudomolecule *vs*. 43.0 ± 20.6 CCS contigs/pseudomolecule)(Figure 2B). Remarkably, in total across the three genomes, fifteen pseudomolecules were reconstructed from a single contig. Haplotype switches were identified based on sequence similarity of protein-coding sequences. Gene loci sequences of each rootstock were aligned against each others using minimap2 v.2.17-r941^20^ and the parameter “-x map-hifi”. Alignments with the highest coverage and identity were used to assign common species parentage and to detect haplotype switches along pseudomolecules (Figure 3A). After manual correction of the haplotype switches, a second cycle of HaploFill^13^ was performed using the pseudomolecules derived from the same *Vitis* species as alternative haplotypes to help closing gaps with draft sequences.

**Figure 2:**
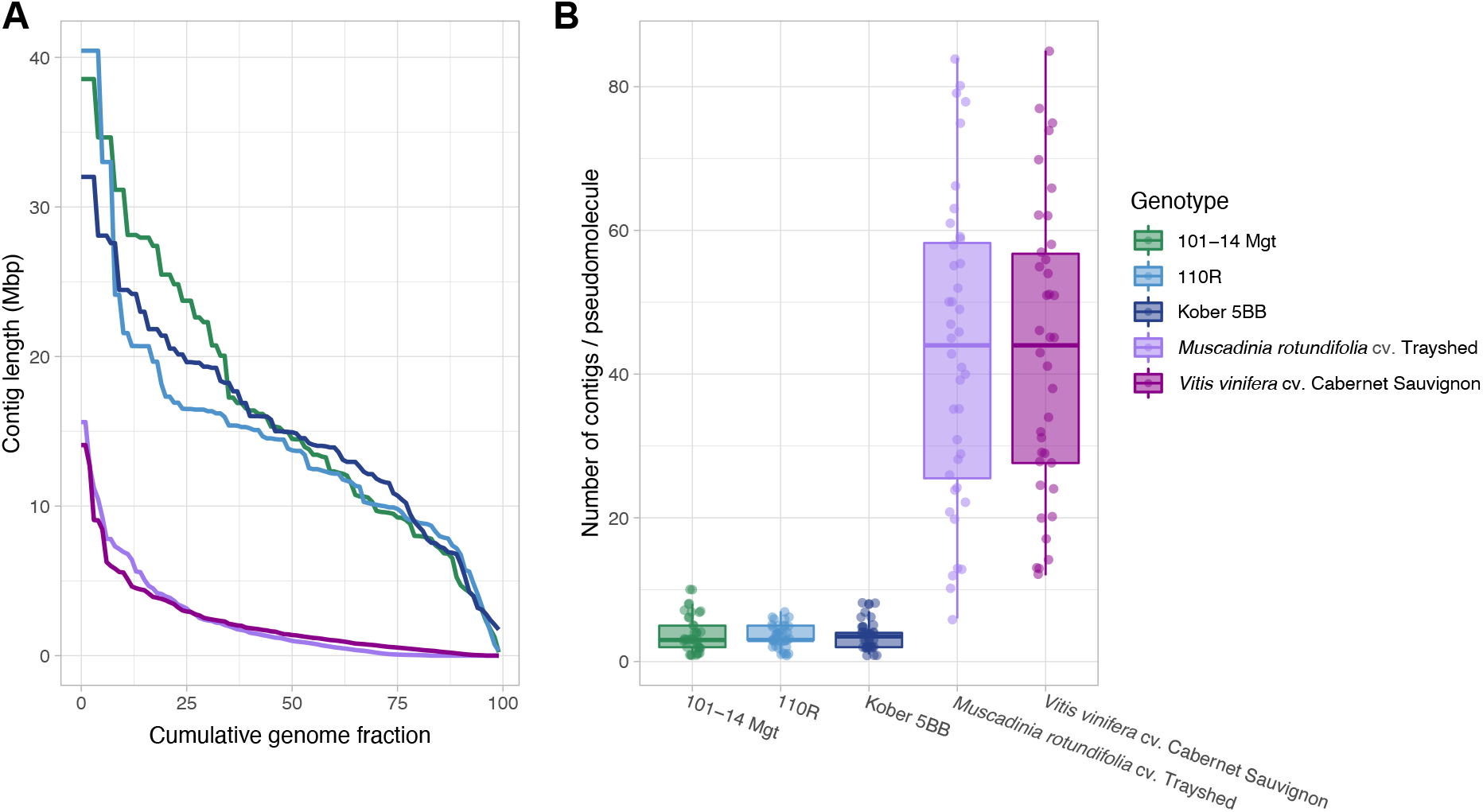
PacBio HiFi sequencing technology substantially improves the contiguity of *Vitis* draft genome assembly. A) Draft assembly fragmentation of 101-14 Mgt, 110R, Kober 5BB represented as distribution of contig NG(x) values. *Muscadindia rotundifolia* cv. Trayshed and *V. vinifera* cv. Cabernet Sauvignon, produced with CCS reads, were included as comparison. The NG(x) value is defined as the sequence length of the shortest contig necessary to achieve, cumulatively, a given fraction (x) of the expected diploid genome length (1 Gbp) when sequences are sorted from the longest to the shortest. Diploid assemblies produced with PacBio HiFi reads (101-14 Mgt, 110R, and Kober 5BB) resulted in a much more contiguous draft genome assembly compared to other grape genomes assembled with older long-read sequencing technologies despite a lower X-Fold coverage employed (PacBio Sequel CLR reads for *M. rotundifolia* 140x X-Fold coverage^19,21^; PacBio RSII CLR reads for Cabernet Sauvignon, 115X X-Fold coverage^14^) B) Distribution of the number of contig scaffolded into complete pseudomolecules . The sustantially lower fragmentation of the draft assemblies generated using PacBio HiFi reads (101-14 Mgt, 110R, and Kober 5BB) resulted on average in a 15x smaller number of contigs necessary to build a pseudomolecule.

**Figure 3:**
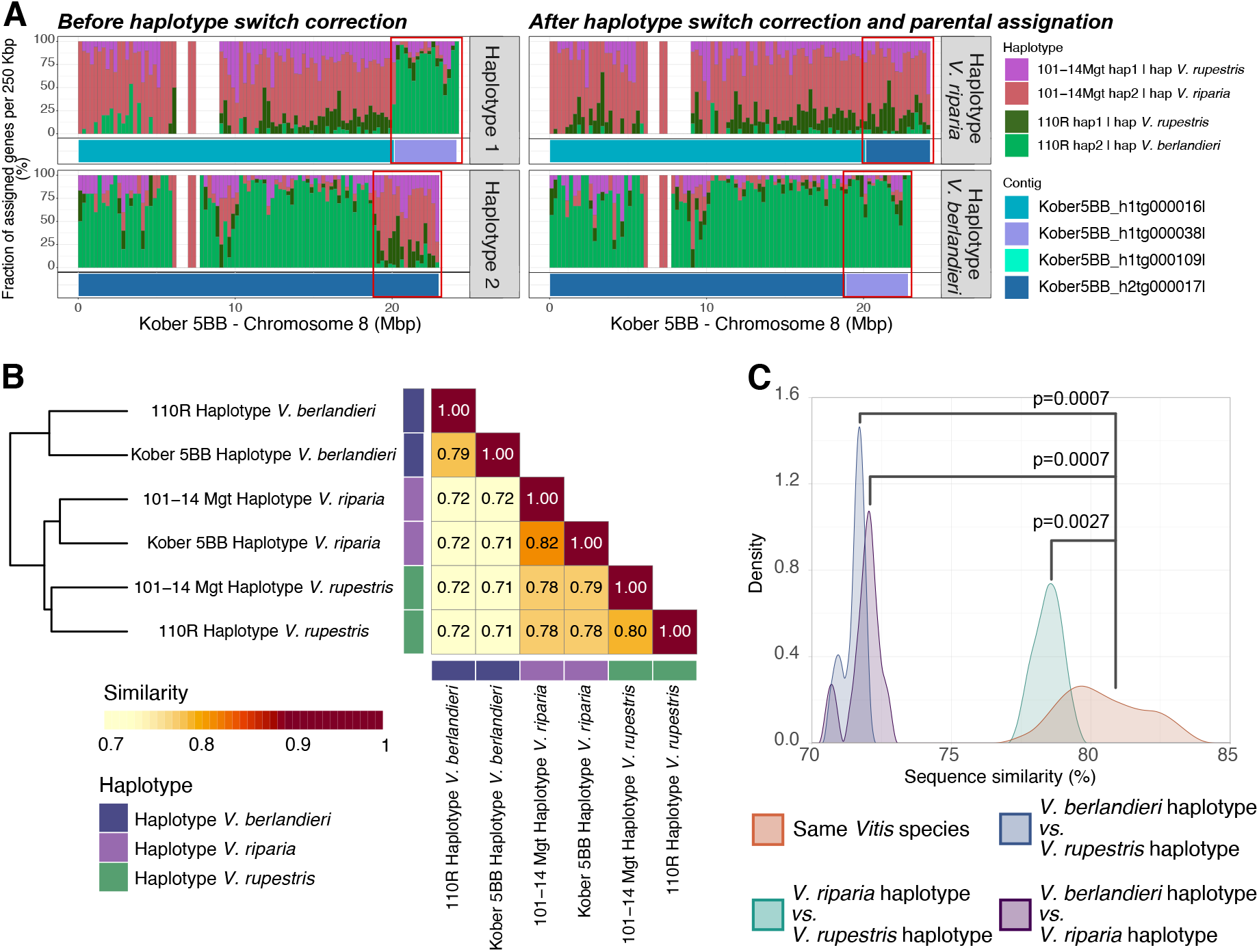
Haplotyping based on intraspecific sequence similarity. Shared parental species information was used to assign each haplotype to either *V. riparia, V. rupestris*, or *Vitis berlandieri* based on sequence similarity. This allowed to resolved assembly errors (i.e. haplotype switches). A) Example of an haplotype switch found on chromosome 8 of Kober 5BB (left panel). After scaffolding of the pseudomolecules, an haplotype switch was observed at the end of chromosome 8 of Kober 5BB. The genes in the contig Kober5BB_h1tg000016l on haplotype 1 were highly similar to the genes located in 101-14 Mgt haplotype 2 (red), suggesting that Kober5BB_h1tg000016l derived from *V. riparia*, whereas the genes of Kober5BB_h1tg000038l corresponded to genes in haplotype 2 of chromosome 8 of 110R (light green), suggesting that Kober5BB_h1tg000038l derived from *V. berlandieri*. An opposite pattern was observed on haplotype 2, with the genes of the first 18.9 Mbp of the pseudomolecule similar to the genes of the haplotype 2 of 110R (light green) and the genes from the last 4.2 Mbp similar to the genes of 101-14 Mgt haplotype 2 (red). The haplotype switch was corrected by interchanging the contig Kober5BB_h1tg000038l with the corresponding region in the alternative haplotype, consisting of Kober5BB_h2tg000109l and 4.2 Mbp of Kober5BB_h2tg000017l (right panel). B) Sequence similarity between haplotypes represented as the average percentage of the haploid chromosome set length not affected by structural variants (> 50bp), SNPs or InDels when compared with another haplotype. C) Distribution of the percentage of sequence similarity (as defined in B) between haplotypes derived from the same species and haplotypes derived from different species (Statistical testing was performed with pairwise Wilcoxon rank sum test).

### Gene prediction and repeat annotation

Gene structural annotations were predicted using the procedures described in https://github.com/andreaminio/AnnotationPipeline-EVM_based-DClab^21^. For each rootstock, Iso-Seq data from the corresponding parental species were concatenated with the *de novo* assembled transcripts from RNA-seq reads before generating the gene models. Iso-Seq libraries underwent extraction, demultiplexing and error correction using IsoSeq3 v.3.3.0 protocol (https://github.com/PacificBiosciences/IsoSeq). Low-quality and single isoforms dataset were further polished using LSC v2.0^22^. RNA-seq reads were quality-filtered and adapters were trimmed with Trimmomatic v.0.36 and the options “ILLUMINACLIP:2:30:10 LEADING:7 TRAILING:7 SLIDINGWINDOW:10:20 MINLEN:36“^23^. High-quality RNA-seq reads from each *Vitis* species were assembled with three different protocols: (i) Trinity v.2.6.5^24^ with the “de novo” protocol, (ii) Trinity v.2.6.5^24^ using the “On-genome” protocol, (iii) Stringtie v.1.3.4d^25^ using the reads found to align on the genome sequences with HISAT2 v.2.0.5 and the parameter “--very-sensitive“^26^. Transcript sequences common to the three assembly methods were then pooled with the Iso-Seq reads. Sequence redundancy was reduced using CD-HIT v4.6^27^ with the parameters “cd-hit-est -c 0.99 -g 0 -r 0 -s .70 -aS.99”. Non-redundant transcripts were processed with PASA v.2.3.3^28^ to obtain the final training model sets. Combined with data from public databases, the derived transcript and protein evidences were aligned on the genome assembly using a multi-aligner pipeline including Exonerate v.2.2.0^29^ and Pasa v.2.3.3^28^. To produce the final set of consensus gene models with EvidenceModeler v.1.1.1^30^, *ab initio* predictions were also generated using Augustus v.3.0.3^31^, BUSCO v.3.0.2^32^, GeneMark v.3.47^33^, and SNAP v.2006-07-28^34^. For the repeat annotation, RepeatMasker v.open-4.0.6^35^ was used. To assign a functional annotation to each of these gene models, results from diamond v2.0.13.151^36, 37^ blastp matches on the Refseq plant protein database (https://ftp.ncbi.nlm.nih.gov/refseq/, retrieved January 17th, 2019) and from InterProScan v.5.28-67.0^38^ were parsed through Blast2GO v.4.1.9^39^. A total of 56,768 protein-coding gene loci were annotated in the genome assembly of 110R, 59,807 in Kober 5BB and 72,758 in 101-14 Mgt. On average, 124,991 ± 36,197 protein-coding alternative splicing variants were identified per haplotype. The unplaced sequences were composed of 2,747 ± 2,821 gene loci (Table 1).

### Analysis of colinearity between haplotypes

Colinear gene loci were identified using MCScanX v.11.Nov.2013^40^. Annotated protein-coding sequences of the three rootstocks were aligned against each other using GMAP v.2019-09-12^41^ with the parameters “-B 4 -x 30 –split-output”. Alignments with both identity and coverage greater than 80% were retained. Alignments corresponding to annotated mRNA regions were identified using mapBed from Bedtools v2.29.2^42^ with the parameters “-F 0.75 -f 0.5 -e”. Colinear blocks were then detected with MCScanx_h (MCScanX v.11.Nov.2013^40^) tool using the following parameters “-s 10 -m 5 -w 5”.

### Identification of sequence polymorphisms and structural variants between haplotypes

Pseudomolecule sequences were aligned against each other using nucmer tool from MUMmer4 v.4.0.0.beta5^43^. SNPs and short indels between haplotypes were identified from alignments with show-snps tool (MUMmer4 v.4.0.0.beta5^43^) with parameters “-Clr-x” and longer structural variants with show-diff tool (MUMmer4 v.4.0.0.beta5^43^) with default parameters.

## Data Records

Sequencing data were deposited at NCBI under BioProject number PRJNA858084. Genome assemblies, gene annotation and repeat annotation files are available at EMBL-EBI under BioProject number PRJEB55013, at Zenodo under the DOI 10.5281/zenodo.6824323, and at http://www.grapegenomics.com. A genome browser and a blast tool are available for each rootstock at http://www.grapegenomics.com.

## Technical Validation

The genome assemblies were evaluated for completeness of the diploid sequence and gene content, and for correct haplotype phasing. The average size of each set of 19 pseudomolecules was 494.5 ± 5.5 Mbp (diploid genome size: 1,015.0 ± 7.9 Mbp), which is close to the length of the parental haploid genome size estimated by flow cytometry (499.3 ± 37.3 Mbp^44^) suggesting that the three genomes were entirely assembled. Only 36.1 Mbp (3.5%), 19.9 Mbp (2.0%), and 23.3 Mbp (2.3%) of the draft sequences could not be placed into any pseudomolecules of 101-14 Mgt, 110R, and Kober 5BB genomes, respectively. The unplaced sequences were mostly composed of repeats (68.0% ± 12.3%). These results are comparable with the latest release of the *V. vinifera* PN40024 reference haploid genome assembly, for which the location of 27.4 Mbp (5.6%) remains undetermined^45^.

Each set of 19 pseudomolecules was evaluated for gene space completeness using both conserved single-copy orthologs of plant genes (BUSCOs) and the single-copy gene content of *Vitis vinifera* PN40024. Complete copies of 98.1 ± 0.14% of the BUSCO models were found in each set of pseudomolecules (Supplemental table 1). Similarly, almost all of the single-copy genes of PN40024 aligned to each set of pseudomolecules (95.01% ± 0.3%). The gene space present in the unplaced sequences was limited to 0.69 ± 0.8% of the BUSCO models and 1.79 ± 0.8% of the PN40024 genes. The completeness of the gene space is another strong evidence that the assemblies are a complete representation of the diploid genomes of the three rootstocks.

Using the pedigree information of each rootstock (Figure 1B), we assigned each pseudomolecule to its parental *Vitis* species, i.e. either *V. riparia, V. rupestris*, or *Vitis berlandieri*. For each pseudomolecule, we identified the three pairs of haplotypes having the highest gene sequence similarity and assigned them to the shared parental *Vitis* species. This allowed us to manually detect and correct the phasing errors (i.e. haplotype switches) introduced during the assembly of the draft sequences or the scaffolding of the pseudomolecules (Figure 3A). Whole-sequence comparison of the six haplotypes of each pseudomolecule showed that the haplotypes assigned to the same *Vitis* species were more similar (80.5% ± 1.4% identity) than those that do not share the same species (74.0% ± 3.3% identity; pvalue = 0.0003, W = 142, n = 30 unpaired Wilcoxon rank sum test; Figure 3B & C). These results suggest that the haplotypes of the three rootstock genomes were correctly phased. Despite the variable levels of sequence polymorphism, pseudomolecules of the three rootstock genomes were highly colinear regardless of their species of origin. When considering both gene sequence similarity, gene order, and physical location, 73.1% ± 3.5% of the protein-coding loci were found in at least one colinear block when comparing haplotypes with shared parental origin, and 71.5% ± 3.5% between haplotypes of different species (Supplemental figure 2). Overall, an average of 82.4% ± 2.6% of the genomic sequences are covered by colinear blocks (Supplemental figure 3), which reflects a remarkable conservation of chromosome structure among these *Vitis* species.

## Code availability

The pipeline used for gene structural and functional annotation is available in details at https://github.com/andreaminio/AnnotationPipeline-EVM_based-DClab

## Acknowledgements

The RNAseq data of *V. rupestris* were kindly provided by Dr. Jason Londo, Cornell University. This work was funded by the NSF grant #1741627 and partially supported by funds to D.C. from the Louis P. Martini Endowment in Viticulture.

## Author contributions statement

A.M., N.C., and D.C. conceived the work. A.M. conducted the bioinformatic analyses. R.F-B. performed all the wet-lab activities associated with the project. A.M., N.C., M.M., D.C wrote the manuscript.

## 1 Supplemental materials

**Supplemental figure 1:**
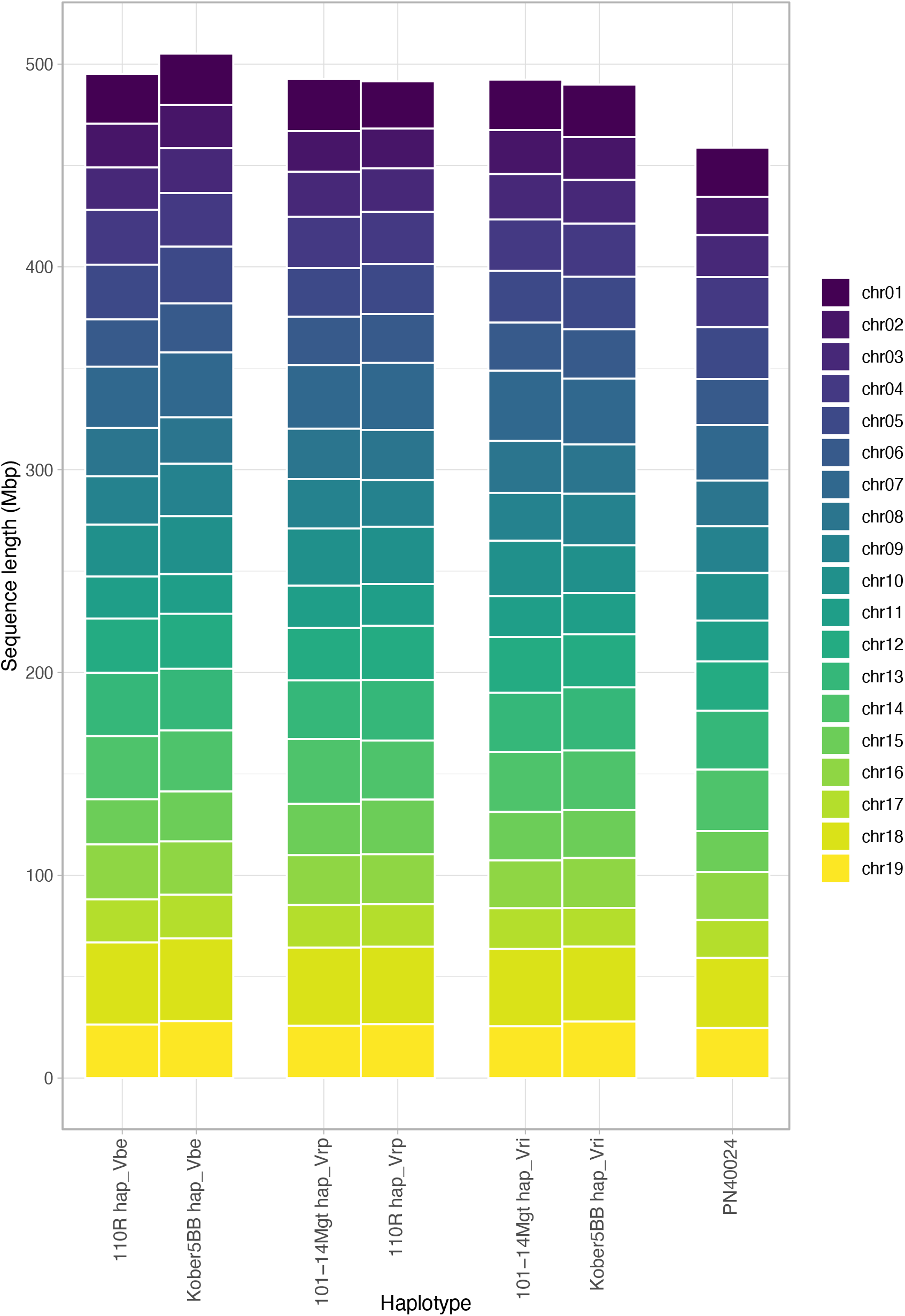
Reconstructed pseudomolecules size comparison. Size of the reconstructed pseudomolecules for each haplotype of the three rootstocks and *V. vinifera* PN40024 genome.

**Supplemental table 1:**
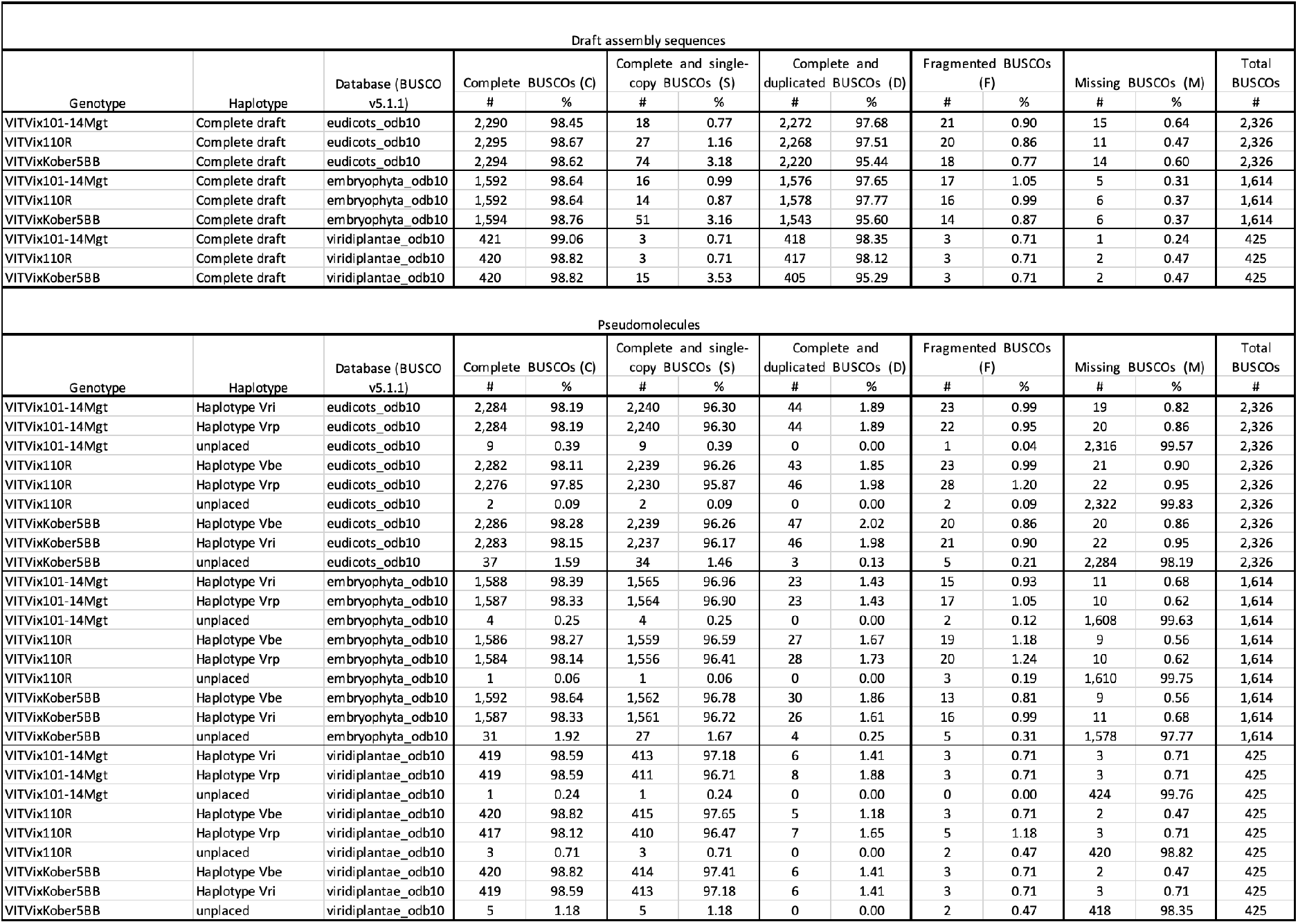
Results of BUSCO search. Results of BUSCO model search in the the assembled sequences for the three rootstocks, before and after pseudomolecule reconstructions using several conserved gene model datasets.

**Supplemental figure 2:**
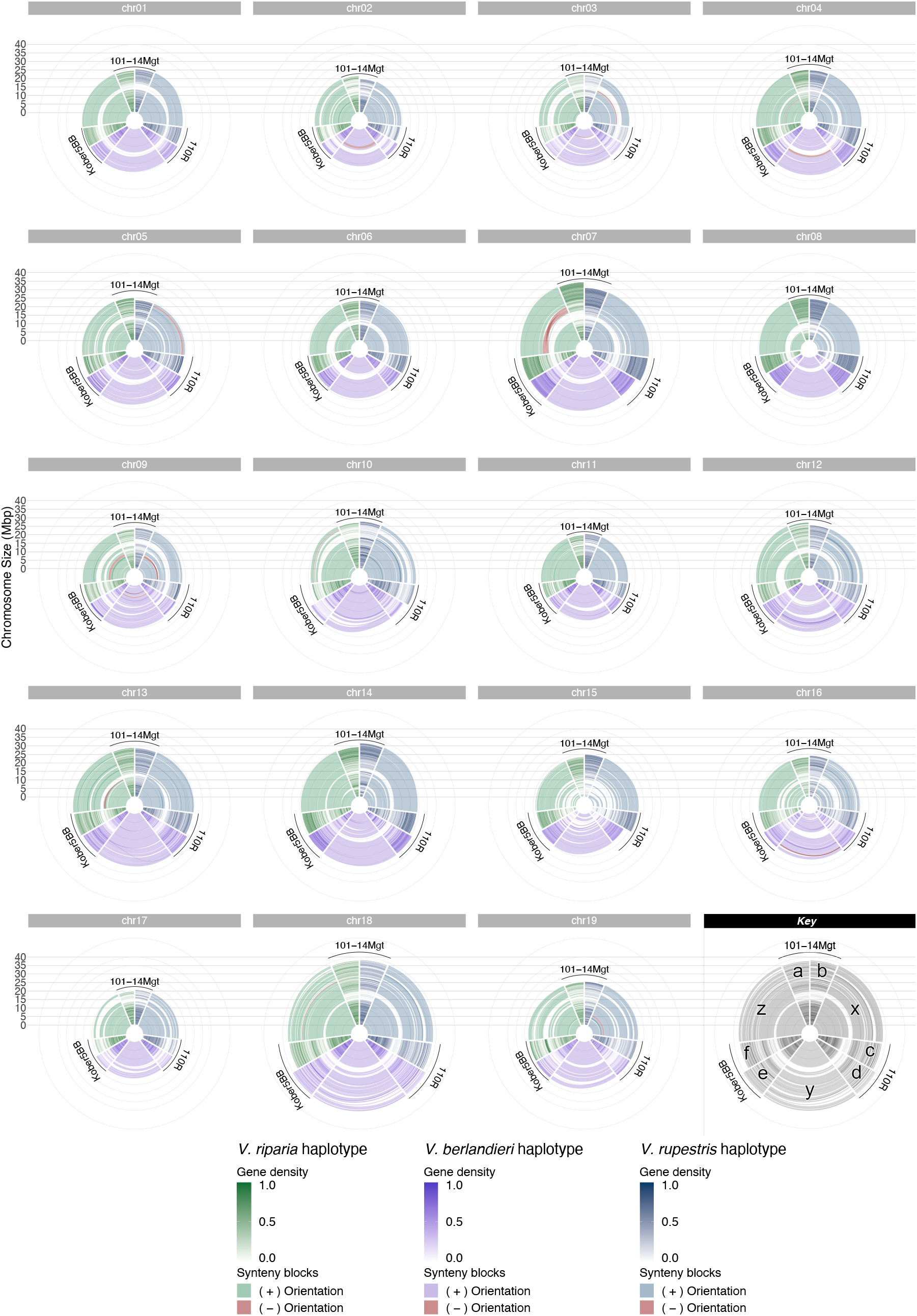
Three-way synteny between 101-14 Mgt, 110R, and Kober 5BB rootstocks highlights the colinearity between siblings. Gene density along 101-14 Mgt haplotypes inherited from *V. riparia* (a) and *V. rupestris* (b), 110R haplotypes from *V. rupestris* (c) and *V. berlandieri* (d), Kober 5BB haplotypes from *V. berlandieri*(e) and *V. riparia* (f). Syntenic regions between the *V. rupestris* haplotypes of 101-14 Mgt and Kober 5BB (x), the *V. berlandieri* haplotypes of 110R and Kober 5BB (y), and the *V. riparia* haplotypes of 101-14 Mgt and 110R (z).

**Supplemental figure 3:**
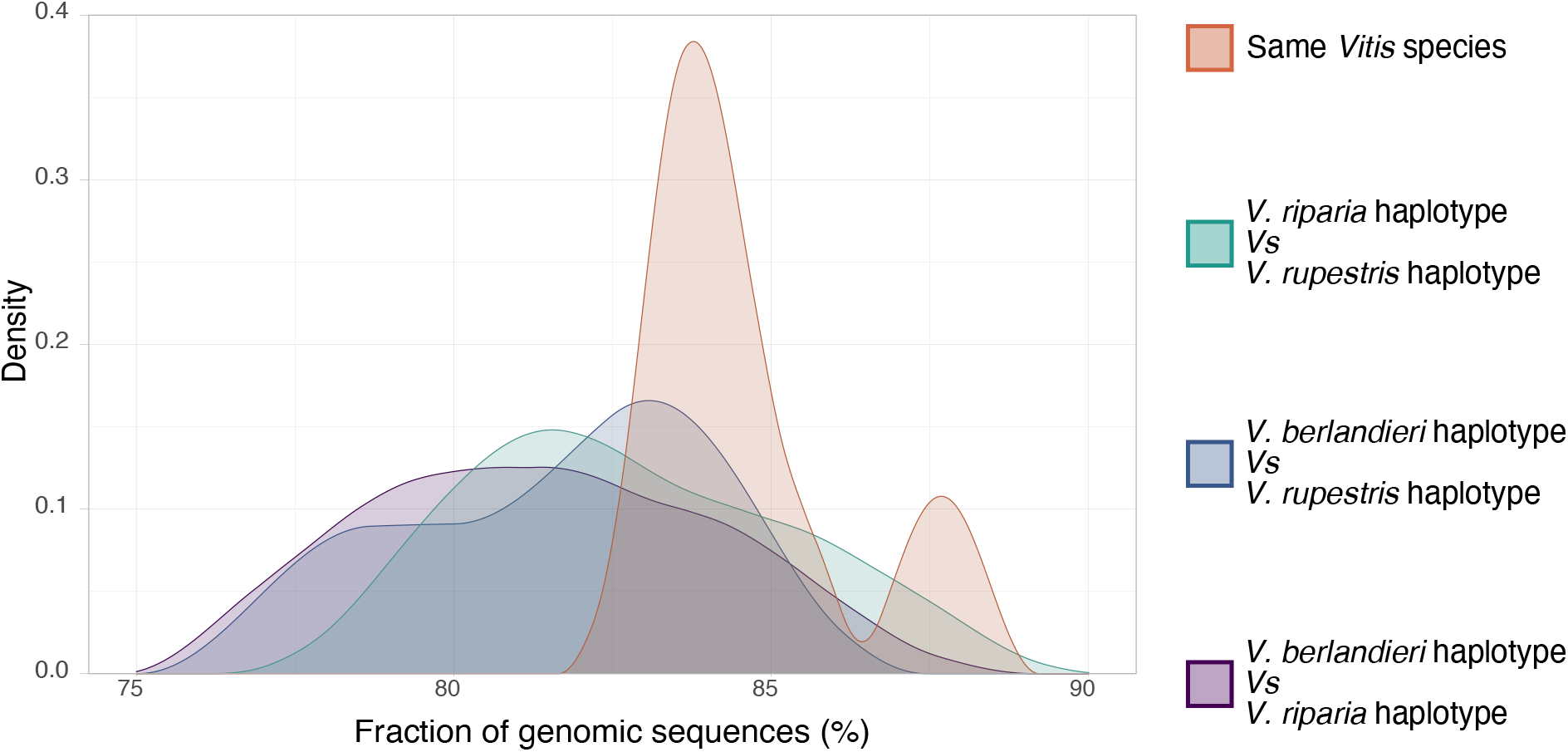
Colinearity between haplotypes. Fraction of the genomic sequences in colinear block of genes between each pair of haplotypes. In average, 82.4% ± 2.6% of the genomic sequences are comprised inside colinear blocks, with very little differences between haplotypes assigned to same the *Vitis* species or to distinct ones.

